# A direct comparison of four high risk human papilloma virus tests versus the cobas test for detecting cervical intraepithelial neoplasia and cervical cancer

**DOI:** 10.1101/314435

**Authors:** Peng Xue, Li-Li Gao, Jian Yin, Li-Li Han, Jing Zhao, Li Li, Samuel Seery, Xue-Yan Han, Ting-yuan Li, Yu Jiang, Jie Shen, Wen Chen

## Abstract

This study is to evaluate performances and genotyping capabilities of four human papilloma virus (HR-HPV) tests based on real-time polymerase chain reaction (PCR) technology platforms compared with the cobas test. Discordant results were further analyzed using INNO-LiPA HPV genotyping test, the gold standard laboratory test to determine presence and type of HPV infection. Over 200 samples from Hospital patients were collected and analyzed using five HR-HPV tests. Women with positive test results were referred directly to colposcopy. If a positive result was returned, biopsies were administered for pathological classification. Clinical performances and genotyping capabilities between the four HR-HPV and cobas tests were compared and contrasted. High levels of agreement were observed, though all HR-HPV tests presented discrepancies compared with the cobas test. Cervical intraepithelial neoplasia Grade 2 or higher lesions (CIN2+) was set as the threshold, and all five tests performed with equally high sensitivity. Lower levels of specificity were observed across all five tests. Results suggest the four HR-HPV tests analyzed are as effective as the cobas test in genotyping capacities and diagnosing CIN. Therefore, these test kits should be used for HPV screening, especially in developing nations because they are cost effective and reliable. Minor discrepancies between tests are generally unavoidable though this may add complexity to the clinical decision-making process. As such, we recommend that efforts be made to standardize HPV genotyping tests as well as to optimize clinical sensitivity and specificity. Focusing on these issues will drive the development of HPV detection techniques, therefore save lives.

## INTRODUCTION

Cervical cancer is associated with a substantial burden of disease and is the cause of a substantial number of deaths among women in developing countries (1). Emerging wealth evidence confirms that persistent infection with high risk human papillomavirus (HR-HPV) is associated with more than 99% of all cervical cancers (2, 3). In particular, HPV 16 and 18 are known to be the most common HPV types, leading to an estimated 70% of all cervical cancers (4, 5).

Accordingly, it has been proposed that HPV detection methods as a alone screening tool for the detection of high-grade cervical intraepithelial neoplasia and cervical cancer (6). The Hybrid Capture 2 (HC2) test is based on a signal-amplified hybridization method and has received approval from the Food and Drug Administration (FDA) yet (7-9), this test can only determine whether HPV infection is present, it neither determines specific genotypes, such as HPV 16 and 18, nor is it capable of identifying between single and multiple HPV infections.

Recently, various tests based on real time polymerase chain reaction (PCR) have been developed. Compared with traditional HC2 tests, the real-time PCR method has several the advantages, such as; convenience of use, high throughput, less time and lower cost. The most frequently administered HPV genotyping test in trials is the cobas HPV test (Roche Systems Inc., Branchburg, NJ, USA). The advantage of the cobas test is that it is a fully automated real-time PCR DNA amplification test which has been approved for screening by the FDA in 2014 (10). The cobas test is initially developed with the clinical cut-off values (11) and this enhances the level of specificity thereby maximizing the predictive value of oncogenic risk of CIN2+. Unfortunately, the cobas test is not flawless, requiring access to highly specialized, bulky instrumentation for sample pretreatment and detection purposes. The cobas HPV system weighs over 150 kg and is 166 cm wide (12). Each detection cost is also comparatively high at $35+ or more per test. These issues make the cobas HPV test impractical and inhibitive for developing nations with less developed infrastructures.

Recently in China, numerous commercially available HR-HPV tests based on real-time PCR have been available in hospitals and laboratories. Unfortunately, prior to 2015 China did not regulate HPV testing around clinical sensitivity or specificity thresholds, and only recently have researchers actually investigated the performances of commonly used HR-HPV tests. Therefore, it would seem necessary to conduct a more comprehensive investigation into the most accessible HPV test kits in order to promote best screening practice for health services in developing countries.

This study focused on four widely used HR-HPV test kits i.e. Tellgen, HybriBio, Liferiver and Sansure, all of which are based on real-time PCR technology. Over 200 samples were collected to appraise and compare the efficacy of each test against the cobas HPV test for detecting HR-HPV DNA. Any discrepant genotyping outcomes were compared with HPV genotyping using INNO-LiPA HPV test in order to determine levels of agreement with the gold standard laboratory diagnostic tool. Levels of sensitivity and specificity HR-HPV test used for diagnosing cervical intraepithelial neoplasia (CIN) and cervical cancer were then analyzed and contrasted. The overarching aim was to determine whether the more cost effective and more easily administered HR-HPV tests can be used for national screening campaigns in developing countries, like China.

## MATERIALS AND METHODS

### Study design and patients

In order to evaluate genotyping capacities and clinical performance of the four HR-HPV tests, we collected a total of 214 cytology samples with cervical lesions results from December 2016 to April 2017. Samples were originally taken from 214 women aged 23 to 65 years whom had visited Peking University First Hospital, for routine examination. Cytology samples collected were transferred into PreservCyt solution (Hologic Inc., Bedford, MA) and then stored at 4°C for testing.

HPV tests were performed using cobas (Roche Molecular Systems Inc., Roche, Shanghai, China), Tellgen (Nucleic Acid Detection Kit for HPV and 16/18 genotyping, Tellgen, Shanghai, China), HybriBio (14 HR-HPV with 16/18 Genotyping Real-time PCR Kit, HybriBio, Guangdong, China), Liferiver (HPV Genotyping Real time PCR Kit, Liferiver, Shanghai, China), and Sansure (HR-HPV DNA Fluorescence Diagnostic Kit, Sansure, Hunan, China) sequentially across specimens. Any discrepant genotyping outcomes between tests were compared using INNO-LiPA HPV test (INNO-LiPA HPV Genotyping Extra, Innogenetics, Belgium).

Women with positive test results were referred directly to colposcopy. If colposcopy returned a positive test result, four-quadrant biopsies were taken. If the colposcopy was unable to detect lesions, a random biopsy was obtained at the squamocolumnar junction in that quadrant at 2, 4, 8, or 10 o’clock.

Informed consent was requested and consequently approved by all participants in this study. This study was formally approved by the institutional review boards of the Cancer Hospital, Chinese Academy of Medical Sciences (NO.12-72/606) and National Health and Family Planning Commission of the People’s Republic of China (No.2015071).

### Real-time PCR HPV testing

A sample of 1 mL of liquid cytology was separated for investigation using five real-time PCR HPV tests i.e. cobas, Tellgen, HybriBio, Liferiver and Sansure all of which are based on TaqMan technology and reportedly can detect 14 HR-HPV genotypes (HPV16, 18, 31, 33, 35, 39, 45, 51, 52, 56, 58, 59, 66 and 68). Besides the 14 HPV types, Liferiver and Sansure can also detect HPV82.

The cobas HPV test has a differentiating feature; the cobas 4800 system is a highly automated instrument for DNA extraction using Roche HPV DNA kit, PCR amplification on the cobas x480 instrument and detection on the cobas z480 Analyzer. The remaining four HR-HPV tests i.e. Tellgen, HybriBio, Liferiver and Sansure perform part of manual DNA extraction using related HPV DNA kits and PCR amplification with a mixture of multiple probes and detection on the ABI 7500 or SLAN-96P automated analyzer.

The experimental conditions for the five HR-HPV tests follow the guidelines provided within the associated protocols. During each run, both positive and negative controls were included to ensure proper PCR responses were not subjected to carry over contamination. The resulting fluorescence from the reaction is then measured to determine whether HPV is present in the sample.

### INNO-LiPA HPV test

The INNO-LiPA HPV test is based on reverse line hybridization using SPF10 primers (13). Part of the L1 region of the HPV genome is amplified by multiplex PCR, and includes biotin-labeled primer which is denatured and hybridized to strip. It can identify 28 HPV types containing 15 HR-HPV (16, 18, 31, 33, 35, 39, 45, 51, 52, 56, 58, 59, 68, 73 and 82), 3 probable HR-HPV (26, 53, 66) and 10 low-risk HPV (HPV6, 11, 26, 40, 43, 44, 54, 69, 70, 71 and 74). Results are interpreted using direct-vision method or utilizing the analytical software, LIRAS for LiPA HPV.

### Statistical analysis

SAS 9.2 software (SAS Institute Inc., Cary, NC) was used for statistical analysis. The agreement rates and corresponding Kappa coefficients with 95% confidence intervals (CIs) were calculated to estimate the level of agreement between the four HR-HPV tests and the cobas HPV test. The Median score and Mann-Whitney U tests were calculated *p* values for the median four HR-HPV and cobas tests Cycle threshold (Ct) values for concordant vs. discordant positive specimens. Chi square test was used to compare the HPV positive rates, sensitivity, specificity, positive predictive value (PPV) and negative predictive value (NPV). *P* values less than 0.05 (two-sided) were considered statistically significant.

## RESULTS

### Overall HPV DNA positivity with the five HR-HPV tests

Of the 214 cytology samples, 4 cases were considered invalid due to lack of remnant DNA and were thereby excluded. The remaining 210 samples were included for analysis. Table 1 displays the HPV DNA positive results with histopathologic grading. Data demonstrate an overall positive correlation of HPV DNA with the histopathologic grading (*p* < 0.0001), except for the positive rate of HPV 18 due to the relative small number of HPV18 type cases within the sample. In 210 cytology samples, overall HPV positive rates ranged from 50.0% to 53.3%. None of the included tests performed less well in overall HPV positive rate (*p* = 0.964).

**Table 1.**
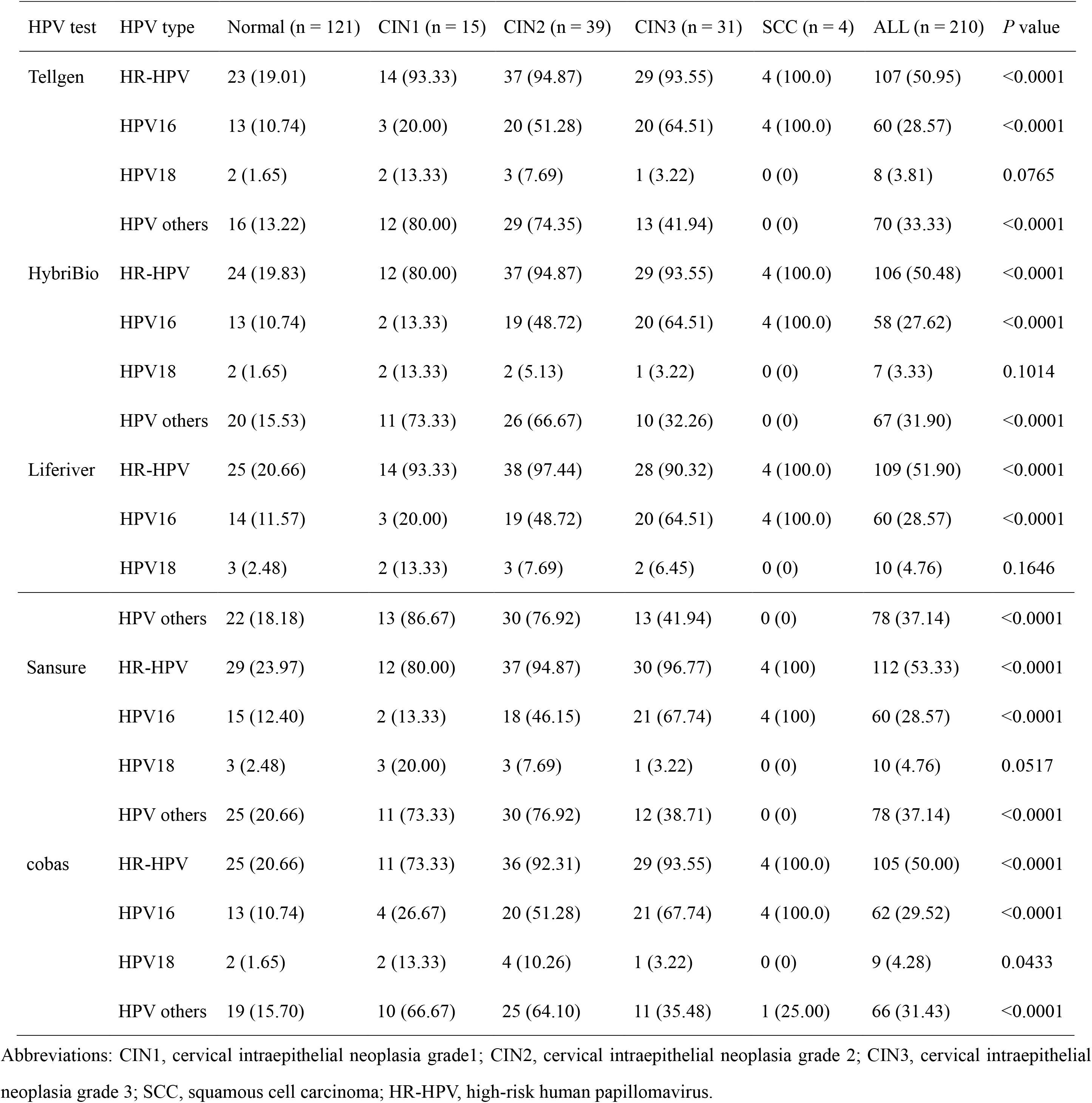
Positivity rates of the five HPV tests according to histopathology classification, n (%)

| HPV test | HPV type | Normal (n = 121) | CIN1 (n = 15) | CIN2 (n = 39) | CIN3 (n = 31) | SCC (n = 4) | ALL (n = 210) | *P* value |
| --- | --- | --- | --- | --- | --- | --- | --- | --- |
| Tellgen | HR-HPV | 23 (19.01) | 14 (93.33) | 37 (94.87) | 29 (93.55) | 4 (100.0) | 107 (50.95) | <0.0001 |
|  | HPV16 | 13 (10.74) | 3 (20.00) | 20 (51.28) | 20 (64.51) | 4 (100.0) | 60 (28.57) | <0.0001 |
|  | HPV18 | 2 (1.65) | 2 (13.33) | 3 (7.69) | 1 (3.22) | 0 (0) | 8 (3.81) | 0.0765 |
|  | HPV others | 16 (13.22) | 12 (80.00) | 29 (74.35) | 13 (41.94) | 0 (0) | 70 (33.33) | <0.0001 |
| HybriBio | HR-HPV | 24 (19.83) | 12 (80.00) | 37 (94.87) | 29 (93.55) | 4 (100.0) | 106 (50.48) | <0.0001 |
|  | HPV16 | 13 (10.74) | 2 (13.33) | 19 (48.72) | 20 (64.51) | 4 (100.0) | 58 (27.62) | <0.0001 |
|  | HPV18 | 2 (1.65) | 2 (13.33) | 2 (5.13) | 1 (3.22) | 0 (0) | 7 (3.33) | 0.1014 |
|  | HPV others | 20 (15.53) | 11 (73.33) | 26 (66.67) | 10 (32.26) | 0 (0) | 67 (31.90) | <0.0001 |
| Liferiver | HR-HPV | 25 (20.66) | 14 (93.33) | 38 (97.44) | 28 (90.32) | 4 (100.0) | 109 (51.90) | <0.0001 |
|  | HPV16 | 14 (11.57) | 3 (20.00) | 19 (48.72) | 20 (64.51) | 4 (100.0) | 60 (28.57) | <0.0001 |
|  | HPV18 | 3 (2.48) | 2 (13.33) | 3 (7.69) | 2 (6.45) | 0 (0) | 10 (4.76) | 0.1646 |
|  | HPV others | 22 (18.18) | 13 (86.67) | 30 (76.92) | 13 (41.94) | 0 (0) | 78 (37.14) | <0.0001 |
| Sansure | HR-HPV | 29 (23.97) | 12 (80.00) | 37 (94.87) | 30 (96.77) | 4 (100) | 112 (53.33) | <0.0001 |
|  | HPV16 | 15 (12.40) | 2 (13.33) | 18 (46.15) | 21 (67.74) | 4 (100) | 60 (28.57) | <0.0001 |
|  | HPV18 | 3 (2.48) | 3 (20.00) | 3 (7.69) | 1 (3.22) | 0 (0) | 10 (4.76) | 0.0517 |
|  | HPV others | 25 (20.66) | 11 (73.33) | 30 (76.92) | 12 (38.71) | 0 (0) | 78 (37.14) | <0.0001 |
| cobas | HR-HPV | 25 (20.66) | 11 (73.33) | 36 (92.31) | 29 (93.55) | 4 (100.0) | 105 (50.00) | <0.0001 |
|  | HPV16 | 13 (10.74) | 4 (26.67) | 20 (51.28) | 21 (67.74) | 4 (100.0) | 62 (29.52) | <0.0001 |
|  | HPV18 | 2 (1.65) | 2 (13.33) | 4 (10.26) | 1 (3.22) | 0 (0) | 9 (4.28) | 0.0433 |
|  | HPV others | 19 (15.70) | 10 (66.67) | 25 (64.10) | 11 (35.48) | 1 (25.00) | 66 (31.43) | <0.0001 |

Abbreviations: CIN1, cervical intraepithelial neoplasia grade1; CIN2, cervical intraepithelial neoplasia grade 2; CIN3, cervical intraepithelial neoplasia grade 3; SCC, squamous cell carcinoma; HR-HPV, high-risk human papillomavirus.

### Agreement among tests

Table 2 displays independent levels of agreement for each of the four HR-HPV tests compared with the cobas test. The four HR-HPV tests, compared with the cobas test demonstrated a high level of agreement with 95.24%, 95.71%, 95.24% and 94.76% of all samples (kappa = 0.905, 0.914, 0.905 and 0.895, respectively). In cases infected with HPV 16, levels of agreement in the four HR-HPV tests against those analyzed using the cobas HPV test were 96.19%, 98.10%, 98.10% and 95.24% (kappa = 0.908, 0.953, 0.954 and 0.885, respectively). In cases infected with HPV 18, agreement with the cobas HPV test were 99.52%, 98.10%, 98.57% and 98.57% (kappa = 0.939, 0.740, 0.835 and 0.835, respectively). HR-HPVs other than HPV types 16 and 18 were also analyzed, and again performed with equally high levels of agreement compared with those of the cobas HPV test with 90.48%, 91.90%, 89.52% and 87.62% (kappa = 0.783, 0.813, 0.768 and 0.726, respectively). 22 discrepancies were eventually resolved using the INNO-LiPA HPV test (see Table 4). 10 cases were negative while LiPA system identified 12 additional positive cases. Histopathologic analysis revealed 6 cases were CIN2+ and 16 were < CIN2. Median Ct values of concordance and discordance in the four HR-HPV tests and cobas test results are presented in Table 3. All concordant cases between the four HR-HPV as well as the cobas HPV tests were significantly lower median cobas Ct values compared to those discordant cases (*p* < 0.001).

**Table 2.**
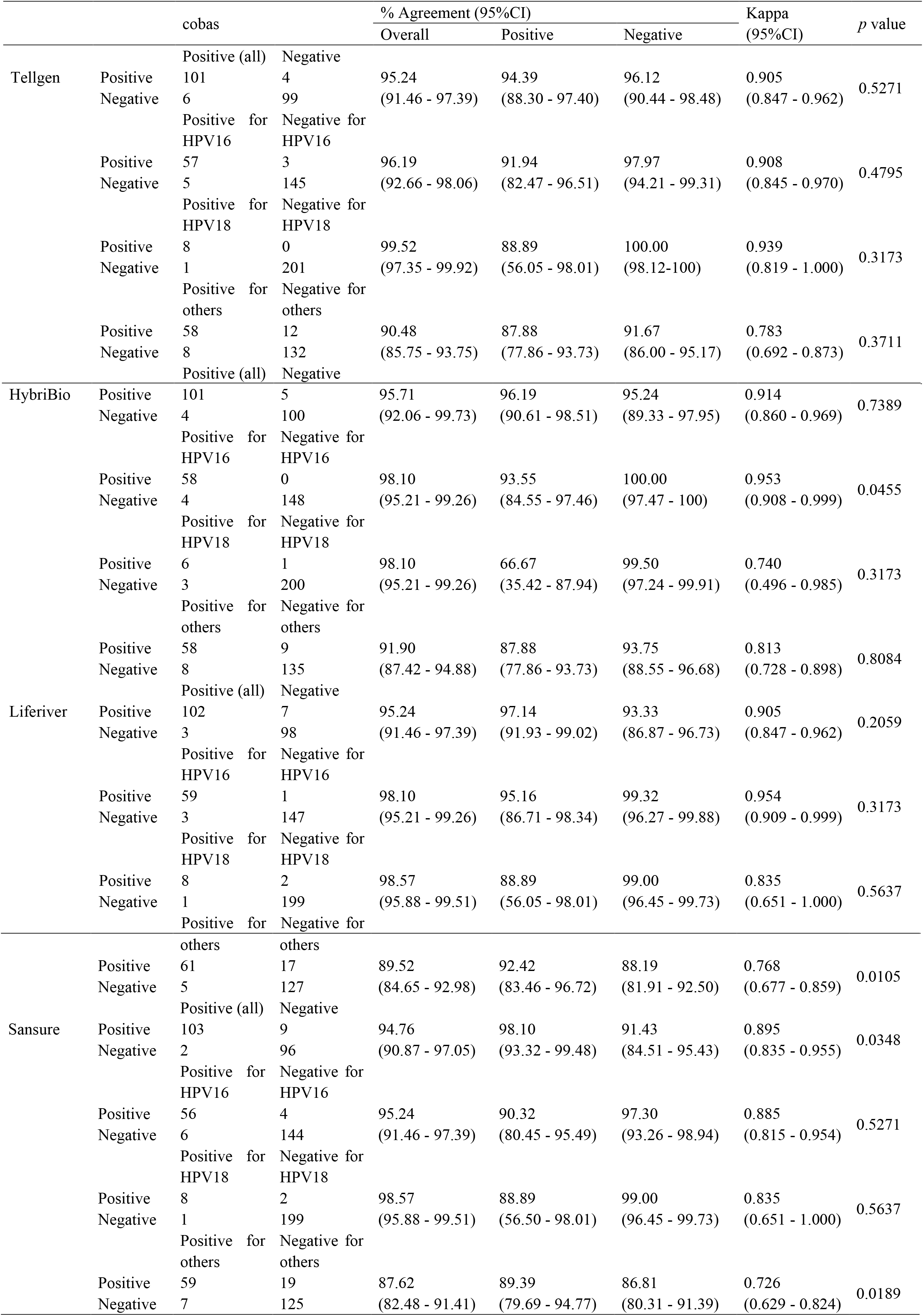
Concordance rates between the results of the four HPV tests and the cobas test

**Table 3.**
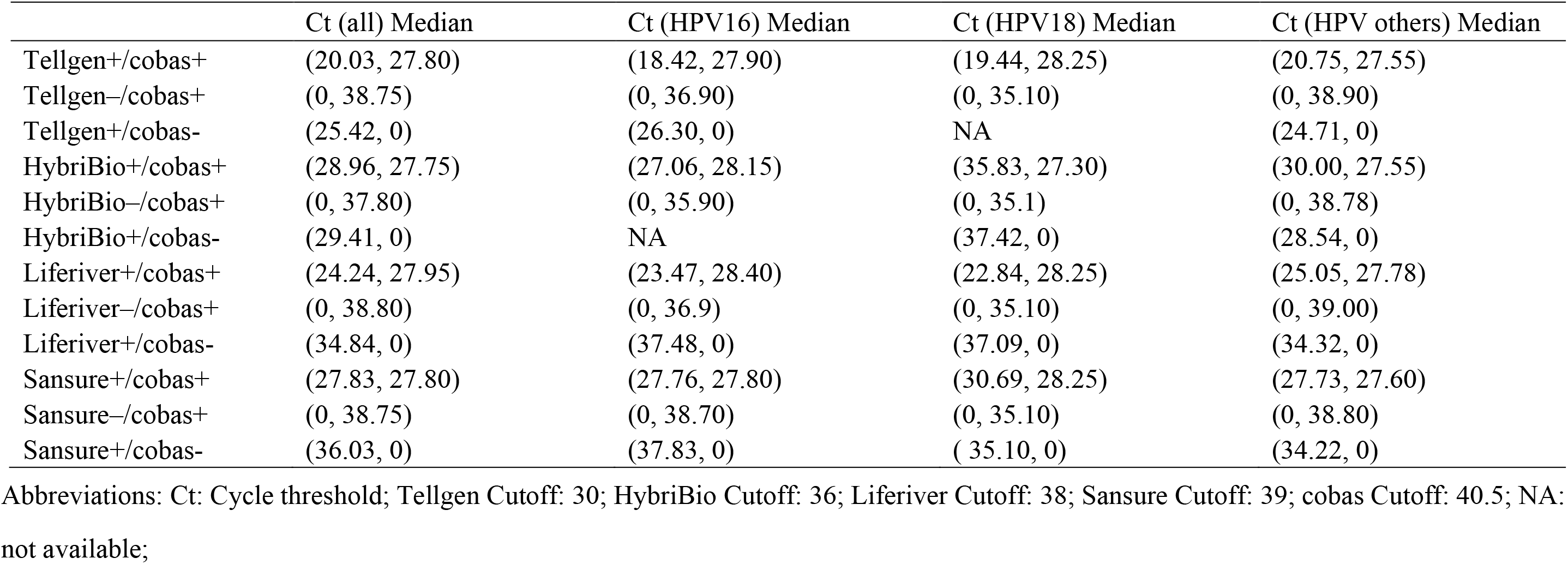
Ct values for concordant and discordant four HPV tests and the cobas test results.

**Table 4.** Discordant results between the four HPV tests and cobas HPV test for HR-HPV compared to histopathology and NNO-LiPA

| Subject ID | Result by:<br>histopathology | Tellgen | HybriBio | Liferiver | Sansure | cobas | INNO-LiPA | Test result interpretation |
| --- | --- | --- | --- | --- | --- | --- | --- | --- |
| A004 | Normal | <u>POS</u> | NEG | <u>POS</u> | <u>POS</u> | NEG | NGE | False positive (Tellgen, Liferiver and Sansure) |
| A008 | CIN2 | <u>NEG</u> | <u>NEG</u> | POS | <u>NEG</u> | <u>NEG</u> | HPV82 | False negative (Sansure) or untargeted <sup>a</sup> |
| A009 | CIN3 | POS | POS | <u>NEG</u> | POS | POS | HPV33 | False negative (Liferiver) |
| A012 | Normal | POS | POS | POS | <u>NEG</u> | <u>NEG</u> | HPV39,66 | False negative (Sansure and cobas) |
| A020 | CIN3 | POS | <u>NEG</u> | POS | POS | POS | HPV16 | False negative (HybriBio) |
| A040 | CIN2 | POS | POS | POS | POS | <u>NEG</u> | HPV52 | False negative (cobas) |
| A042 | CIN3 | <u>NEG</u> | POS | POS | POS | POS | HPV16 | False negative (Tellgen) |
| A054 | Normal | NEG | NEG | <u>POS</u> | NEG | NEG | NEG | False positive (Liferiver) |
| A064 | CIN3 | POS | POS | <u>NEG</u> | POS | <u>NEG</u> | HPV53 | False negative (Liferiver and cobas) or untargeted |
| A075 | Normal | <u>POS</u> | NEG | NEG | NEG | NEG | NEG | False positive (Tellgen) |
| A100 | Normal | POS | POS | POS | POS | <u>NEG</u> | HPV52 | False negative (cobas) |
| A112 | Normal | NEG | NEG | NEG | NEG | <u>POS</u> | NEG | False positive (cobas) |
| A127 | Normal | NEG | NEG | NEG | <u>POS</u> | NEG | NEG | False positive (Sansure) |
| A128 | Normal | NEG | NEG | <u>POS</u> | NEG | NEG | NEG | False positive (Liferiver) |
| A131 | Normal | NEG | NEG | NEG | <u>POS</u> | NEG | NEG | False positive (Sansure) |
| A132 | Normal | NEG | NEG | NEG | <u>POS</u> | NEG | NEG | False positive (Sansure) |
| A133 | Normal | NEG | NEG | NEG | <u>POS</u> | NEG | NEG | False positive (Sansure) |
| A187 | Normal | NEG | NEG | NEG | NEG | <u>POS</u> | NEG | False positive (cobas) |
| A192 | Normal | POS | <u>NEG</u> | POS | POS | POS | HPV16,56 | False negative (HybriBio) |
| A197 | Normal | NEG | NEG | NEG | <u>POS</u> | NEG | HPV43 | False negative (cobas) |
| A202 | Normal | <u>NEG</u> | POS | POS | POS | POS | HPV18,31,52,59,74 | False negative (Tellgen) |
| A209 | Normal | <u>NEG</u> | POS | <u>NEG</u> | <u>NEG</u> | <u>NEG</u> | HPV18,51 | False negative (Tellgen, Liferive, Sansure and cobas) |

Abbreviations: POS: Positive; NEG: Negative; CIN2: cervical intraepithelial neoplasia grade 2; CIN3: cervical intraepithelial neoplasia grade 3;
a: Lesion caused by HPV genotypes not targeted by cobas or the four HR-HPV tests; Underline: discordant results with LiPA test.

### Clinical performance of the five HR-HPV tests for detection of CIN2+

Clinical performances of each of the HR-HPV tests as well as the cobas HPV test to a reference standard of CIN2+ were analyzed (see Table 5). Data revealed that all tests were similarly sensitive with 94.59%, 94.59%, 94.59%, 95.59% and 93.24%, respectively. However, each of the tests was significantly less specific. HR-HPV tests i.e. Tellgen, HybriBio, Liferiver, Sansure and the cobas HPV test resulted in 72.79%, 73.53%, 71.32%, 69.86% and 73.53%, specificity. Overall, there was no significant difference in either sensitivity (*p* = 0.971) or specificity (*p* = 0.953) across these HR-HPV tests for detecting CIN2+.

**Table 5.** Clinical performance of these HR-HPV tests for detection of CIN2+ in women with positive HPV results.

| Endpoint | HPV test | Sensitivity (95%CI) | Specificity (95%CI) | PPV (95%CI) | NPV (95%CI) |
| --- | --- | --- | --- | --- | --- |
| CIN2+ (n = 74) | Tellgen | 94.59 (86.91 - 97.88) | 72.79 (64.77 - 79.57) | 65.42 (56.02 - 73.76) | 96.12 (90.44 - 98.48) |
|  | HybriBio | 94.59 (86.91 - 97.88) | 73.53 (65.54 - 80.22) | 66.04 (56.60 - 74.35) | 96.15 (90.53 - 98.49) |
|  | Liferiver | 94.59 (86.91 - 97.88) | 71.32 (63.22 - 78.26) | 64.22 (54.88 - 72.59) | 96.04 (90.26 - 98.45) |
|  | Sansure | 95.95 (88.75 - 98.61) | 69.86 (61.68 - 76.93) | 63.39 (54.17 - 71.73) | 96.94 (91.38 - 98.95) |
|  | cobas | 93.24 (85.14 - 97.08) | 73.53 (65.54 - 80.22) | 65.71 (56.23 - 74.09) | 95.24 (89.33 - 97.95) |

Abbreviations: CIN2+: cervical intraepithelial neoplasia grade 2 or higher; CI: confidence interval; PPV: positive predictive value; NPV: negative predictive value.

## DISCUSSION

This study focused on four readily available and widely used HR-HPV test kits i.e. the Tellgen, HybriBio, Liferiver, and the Sansure, all of which are based on real-time PCR technology. Clinical performance and genotyping capabilities were compared with the cobas HPV test. Discrepancies were further compared with HPV genotyping using INNO-LiPA HPV test, the gold standard laboratory diagnostic test. Since approval by the U.S. FDA and validation by numerous studies (14, 15), the cobas HPV test is as effective as the Hybrid Capture 2 HPV test which has become a gold standard screening tool for evaluating the efficacy of the newly developed HPV methods (16). In addition, INNO-LiPA has been widely used in clinical trials focusing on HPV vaccine research for the identification of specific sequences in the L1 region of the HPV genome of 28 HPV types (17).

Considering that the four HR-HPV tests; the Tellgen, HybriBio, Liferiver, and Sansure have become widely available within hospitals and laboratories in China, there were few studies which compare and then verify test performances simultaneously. In this study, by using CIN2+ as a threshold and reference standard, it became possible to compare levels of sensitivity and specificity of all four HR-HPV tests which ultimately performed very similar when compared with the cobas HPV test. Similar HPV-DNA detection rates were also discovered compared with both the cobas HPV test and the INNO-LiPA HPV test. This study ultimately found that these HR-HPV tests and cervical histopathology positively correlate. All HPV tests detected HPV from CIN2+ samples in approximately 90.3% to 100.0% of cases therefore this analysis demonstrates that these cheaper, simpler HR-HPV tests perform equally at detecting HPV infection in cervical lesions.

All four HR-HPV tests demonstrated high levels of agreement with the cobas HPV test for detecting HR-HPV DNA. However, discrepancies detected by these HR-HPV tests in 22 (10.5%) of the 210 samples, thus genotyping using INNO-LiPA was performed to explore potential causes. The cobas-negative/four HR-HPV tests-positive samples resulted in approximately 70% of the all discrepancies, the four HR-HPV-positive results found in half of the cases were not confirmed by LiPA, whereas, the LiPA detection test detected the presence of different low-risk genotypes, thus representing false-positive results though the four HR-HPV tests. These differences may lead to over-referral to colposcopy and potentially false prognostic stress in healthy women.

On the other hand, 7 samples were cobas-positive/four HR-HPV-negative; 5 samples returned positive for HR-HPV using the LiPA system, with 3 cases bearing a CIN2+, which were later identified as HPV 33 and 16. It should be noted that LiPA does not have an established clinical cut off value and it was expected that the positive sample below the critical cut off point would be detected using these HR-HPV tests. Genotype-specific results detected using these HR-HPV tests showed that samples with discordant genotyping results had Ct values significantly closer to the test limits of detection which was consistent with genotyping samples. This indicates that these samples may have contained a lower viral load based on high Ct values and are more likely to produce discordant results across the tests (18). These minor discrepancies were considered unavoidable due to different cut-off values and a lack of standardization. No test is absolutely sensitive or specific and we therefore suggest that cut-off values for HR-HPV tests need to be adjusted for optimization in order to reduce false-positive results and without sacrificing sensitivity in detecting HR-HPV.

This study had some limitations that should be addressed. First of all, this is a preliminary study with a relatively small sample (n = 210). Those results led to a high-quality appraisal of four commonly used HR-HPV tests however due to the small sample size our recommendations remain tentative. Additional research with a large sample size is required to verify these findings. Secondly, the total HR-HPV concordance rates were perhaps overestimated due to the fact that detection methods identify HPV types as a pool rather than each HPV Genotype. Further research is required to analyse test capabilities for each HPV genotype. A more powerful form of sequencing generation could also be used, which may provide more definitive HPV genotype information. Lastly, manual operation of the four HR-HPV tests were time consuming and labor intensive and were therefore more likely to suffer potential sample cross-contamination. We suggest the manufacturer should use an automated DNA extraction system if conditions are available in order to minimize potential cross-contamination as well as to reduce hands-on time.

Ultimately, this study demonstrates that the four included HR-HPV tests have strong levels of agreement and similar clinical performance when compared with the cobas HPV test. Therefore, nations with less developed healthcare systems should consider using one of these test kits for HPV and for national cervical cancer screening. Slight discrepancies between tests are generally unavoidable. It is therefore recommended that HPV genotyping tests should be optimized and standardized around test sensitivity and specificity. This will enhance HPV detection techniques and reduce the burden of cervical cancer in developing regions and countries. Further research with larger samples and comparison with various HPV detection methods is of course required.

## ACKNOWLEDGMENTS

This study was supported by National Natural Science Foundation of China (NO.71373166) The authors have declared that no competing interest exists.

